# Heterozygous gene truncation delineates the human haploinsufficient genome

**DOI:** 10.1101/010611

**Authors:** István Bartha, Antonio Rausell, Paul J McLaren, Pejman Mohammadi, Manuel Tardaguila, Nimisha Chaturvedi, Jacques Fellay, Amalio Telenti

## Abstract

Sequencing projects have identified large numbers of rare stop-gain and frameshift variants in the human genome. As most of these are observed in the heterozygous state, they test a gene’s tolerance to haploinsufficiency and dominant loss of function. We analyzed the distribution of truncating variants across 16,260 protein coding autosomal genes in 11,546 individuals. We observed 39,893 truncating variants affecting 12,062 genes, which significantly differed from an expectation of 12,916 genes under a model of neutral *de novo* mutation (*p*< 10^−4^). Extrapolating this to increasing numbers of sequenced individuals, we estimate that 10.8% of human genes do not tolerate heterozygous truncating variants. An additional 10 to 15% of truncated genes may be rescued by incomplete penetrance or compensatory mutations, or because the truncating variants are of limited functional impact. The study of protein truncating variants delineates the essential genome and, more generally, identifies rare heterozygous variants as an unexplored source of diversity of phenotypic traits and diseases

---

Recent population expansion and limited purifying selection have lead to an abundance of rare human genetic variation ^1–3^ including stop-gain and frameshift mutations. Thus, there is increasing interest in the identification of natural human knockouts ^3–8^ through the cataloguing of homozygous truncations. However, heterozygous truncation can also lead to deleterious functional consequences through haploinsufficiency due to decreased gene dosage, or through a dominant-negative effect ^9,10^. In order to quantify the importance of heterozygous protein truncating variation, we characterized genes showing fewer *de novo* truncations in the general population than expected under a neutral model. We hypothesized that there is a set of genes that cannot tolerate heterozygous protein truncating variants (PTVs) because of early life lethality.

## Results

### Fewer genes carry heterozygous PTVs than expected under neutral evolution

We used stop-gain (nonsense) single nucleotide variants and frameshift (insertions/deletions) variants to assess tolerance to heterozygous PTVs across the human genome. We considered transcripts from 16,260 autosomal protein coding genes annotated by the consensus coding sequence (CCDS) project ^11^, for which *de novo* mutation rate estimates were recently calculated ^12^, and where the number of synonymous variants in sequenced individuals followed expectation (**Online Methods**). The study dataset included 11,546 exomes in which we observed 39,893 rare PTVs (allele frequency < 1%), affecting 12,062 (74.1%) genes.

To test whether there is a subset of genes that are intolerant to heterozygous truncation, we simulated a model of generation of neutral *de novo* PTVs for all genes (i.e. assuming viability of affected individuals). By randomly assigning 39,893 hypothetical stop-gain and frameshift variants to genes according to their *de novo* mutation rate ^12^, we observed that 12,916 out of 16,260 genes (95% CI, 12,805-12,991) would be expected to carry at least one stop-gain or frameshift variant. The expected number of genes is significantly greater than the 12,062 truncated genes observed in the study dataset for the same number of PTVs (6.6% depletion, empirical p-value < 1×10^−4^; **Figure 1A**). The depletion in number of observed truncated genes was greater when severe PTVs, i.e. those predicted to have the greatest functional impact ^13^, were considered (n=10,340 vs. a neutral expectation of 11,821-11,978; 13.1% depletion p < 1×10^−4^). This suggests that a measurable fraction of *de novo* heterozygous stop-gain and frameshift variants are highly deleterious and hence under strong purifying selection. Hereafter we denote that fraction as the haploinsufficient genome (*f*_*hi*_).

**Figure 1.**
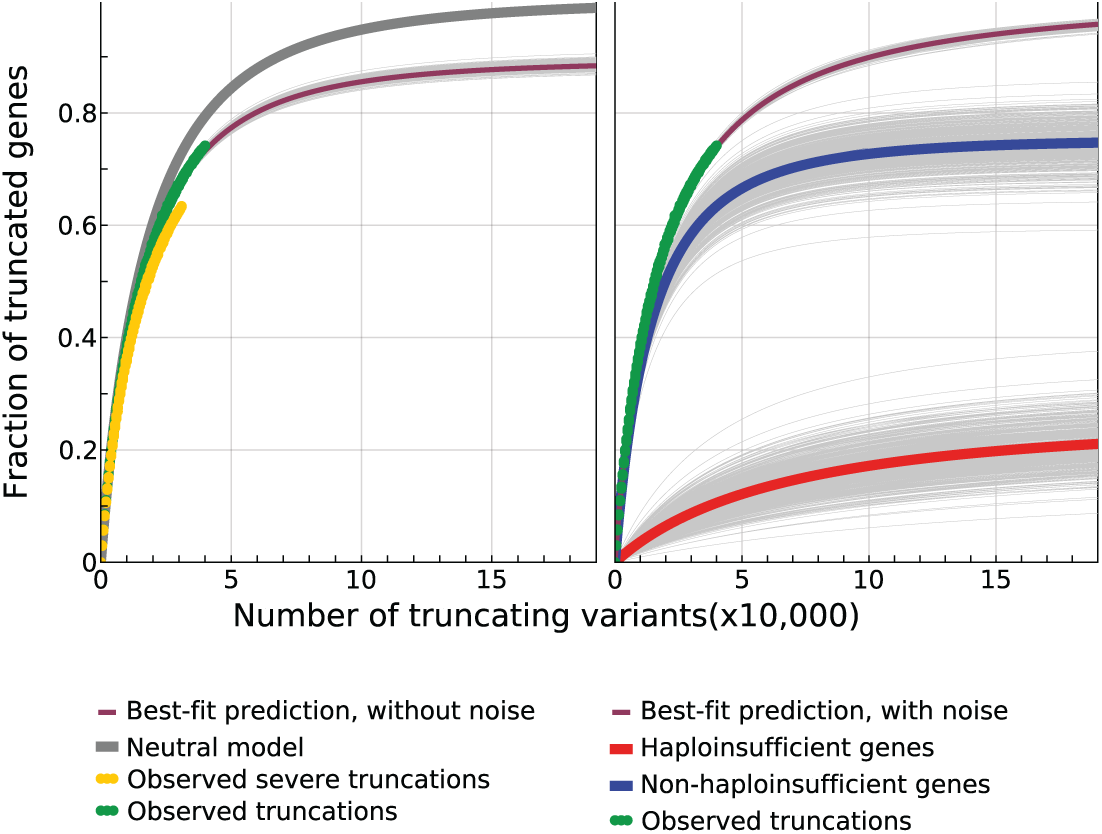
Observed and expected PTVs in the study population. **A**: Fraction of genes with at least one stop-gain or frameshift variant as a function of the number of sampled PTVs. The gray curve shows the expected number of genes under a model of neutral de novo mutation rate 12 representing the null hypothesis (no deleterious effects). The green curve shows the number of genes observed with at least one PTV. The orange curve limits the number of observed genes to those hosting highly damaging variants 13. The purple curve shows the predicted number of genes with at least one PTV under the estimated best-fit parameters under model A (see Online Methods). **B**: Extrapolation of the observed number of genes with at least one PTV assuming a model that includes the possibility of finding PTVs due to biological and technical noise. The purple curve shows the predicted number of genes with at least one PTV under the estimated best-fit parameters, while the green curve shows the observed data. Decomposition of the observed and predicted number of genes with at least one PTV: variants in non-haploinsufficient genes (blue) saturate early; variants found in haploinsufficient genes (red) continue to accumulate PTVs due to the constant contribution of biological and technical noise.

### Characteristics of genes comprising the haploinsufficient genome

We assessed the functional properties of the subset of genes that were not observed to carry PTVs (n=4,198), **Table 1**. These genes were highly conserved, had fewer paralogs, were more likely to be part of protein complexes and were more connected in protein-protein interaction networks than the rest of the genes. Furthermore, they had characteristics of essentiality and haploinsufficiency, and a higher probability of CRISPR-Cas9 editing compromising cell viability. The set of genes not carrying PTVs was enriched in OMIM genes annotated with ‘haploinsufficient’ or ‘dominant negative’ keywords. Non truncated genes were overrepresented in functional categories such as transcription regulation, developmental processes, cell cycle, and nucleic acid metabolism (**Supplementary Table 1)**, in line with earlier characterization of haploinsufficient genes ^14^. Together, these results indicate that a number of basic cellular functions depend on the integrity of coding and expression of both alleles of component genes.

**Table 1.** Characteristics of the subset of genes (n=4,204) observed without PTVs after sequencing 16,260 protein coding autosomal genes in 11,546 individuals. Tests compare genes with and without heterozygous PTVs.

| Annotation | Effect in non-truncated genes | P-value | Test | Data Source |
| --- | --- | --- | --- | --- |
| dN/dS | Lower (conservation) | 1E-295 | Rank-sum test | Ensembl primate genomes<sup>13</sup> |
| Paralog count | Lower | 4E-94 | Poisson regression | Ensembl Biomart |
| Loss of cell viability (CRISPR-Cas9) | Enrichment | 3E-16 | Logistic regression | Shalem et al. 2014<sup>26</sup> |
| Part of a protein complex | Enrichment | 3E-29 | Logistic regression | Gene Ontology term “Protein complex” GO:0043234 |
| Essentiality | Higher | 4E-34 | Logistic regression | OGEE (http://ogeedb.embl.de/) |
| Connectivity in protein-protein interaction network | Higher | 5E-52 | Linear regression | OGEE (http://ogeedb.embl.de/) |
| Predicted haploinsufficiency | Higher | 1E-162 | Linear regression | Huang et al. 2010<sup>10</sup> |
| OMIM ‘haploinsufficient’ and ‘dominant negative’ subset | Enrichment | 5E-12 | Logistic regression | Petrovski et al. 2013<sup>32</sup> |

### Estimating the fraction of genes intolerant to heterozygous stop-gain and frameshift variants

Genes without PTVs in our analysis may be truly part of the haploinsufficient genome or the result of insufficient sample size to detect rare events. Thus, we next sought to estimate the total haploinsufficient fraction (*f*_*hi*_) of the genome in the full population by a modeling approach. Assuming that a fraction *f*_*hi*_ of genes do not carry *de novo* PTVs while the remaining genes do so according to their neutral mutation rates ^12^, *f*_*hi*_ can be estimated by fitting a model to the observed relative distribution of PTVs (relative to the rest of genes; **Online Methods**). This analysis estimates a fraction of the haploinsufficient genome of *f*_*hi*_ = 10.8% (95% CI=9.5-11.7%) of protein coding genes (**Figure 1A**).

Some genes may tolerate PTVs because their functional effects are masked by incomplete penetrance ^15^, by compensatory variants ^16^, or because of a low functional impact of the truncation ^13^. In addition, false positive errors in sequencing and variant calling procedures contribute to the distribution of observed variants ^17–19^. We collectively treated these factors as noise, because they can lead to the observation of a truncated gene in a viable individual without truly probing the general viability of carrying only one functional allele in a given gene. Therefore, we extended our model to allow for the possibility of observing PTVs in the haploinsufficient fraction of the genome by introducing a second parameter representing the number of variants originating from biological noise (incomplete penetrance, compensatory variants and low impact truncation) or technical noise (sequencing or variant calling errors) in genes otherwise intolerant to truncation (**Online Methods**). Using these parameters, the estimated fraction of genes intolerant to PTVs increased to 24.4% (95% CI, 18.3-32.1%, **Figure 1B**).

An important consequence of biological and technical noise is that the apparently truncated fraction of genes does not saturate as a function of the number of observed PTVs, but keeps rising. Our model predicts that after having sequenced 40,000 exomes (representing a sample of approximately 90,000 PTVs) more than 50% of newly identified truncated genes will result from biological and technical noise (**Supplementary Figure 1)** - an important consideration for ongoing sequencing programs and interpretation of resources, such as that of the Exome Aggregation Consortium (ExAC, http://exac.broadinstitute.org). At the sample size of 40,000 exomes, and with 2 to 6% of all observed truncations due to technical errors ^5,6,8^, 400 to 1025 genes intolerant to PTVs will exhibit truncations due to sequencing and variant calling errors. For the same sample size, 2345 to 2549 genes intolerant to PTVs will exhibit truncations due to incomplete penetrance, compensatory variants or low impact truncation.

We next assessed the robustness of these estimates using an alternative approach that models the expected number of PTVs as a function of the observed synonymous coding variants (**Online Methods)**. This model assumes that, in the absence of deleterious consequences, the number of heterozygous PTVs correlates with the number of synonymous variants observed in a gene. This approach resulted in highly similar estimates of *f*_*hi*_ (95% CI 19.7-34.1%) compared to the previous model. Leveraging the latter model, we identified 278 genes (**Supplementary Table 2**) that have higher than 0.99 posterior probability of being intolerant to heterozygous truncation (**Figure 2**). However, there is a continuum of tolerance to heterozygous truncation as depicted in **Figure 2**, with a large number of genes harboring fewer heterozygous PTVs than expected.

**Figure 2.**
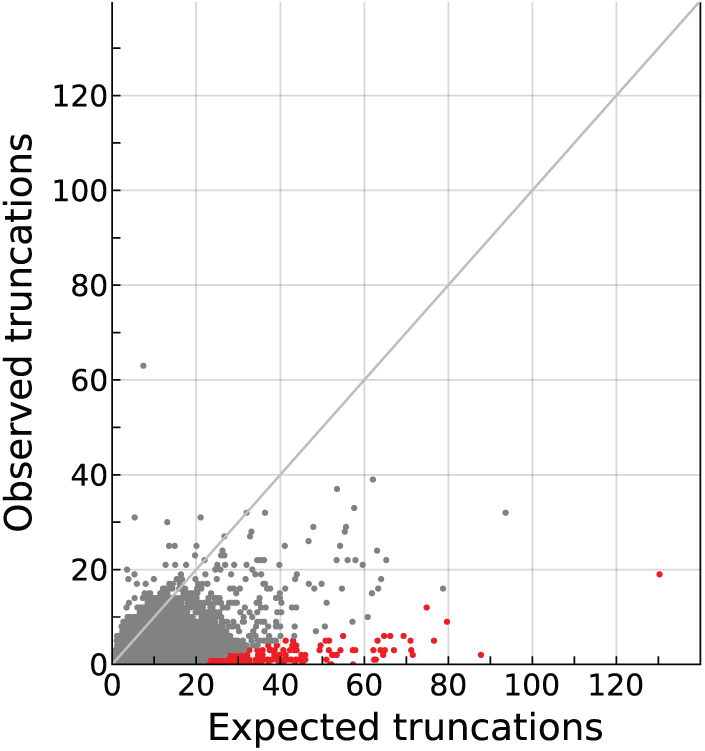
Expected and observed number of PTVs per gene. Each dot in the scatter plot corresponds to a gene. X-axis reflects the expected number of PTVs for each gene according to a model of neutral variation based on synonymous variants (Model B, see Online Methods) while on Y-axis indicates observed number of PTVs in the study dataset. Genes intolerant to heterozygous PTVs with a posterior probability of ≥ 0.99 are colored in red. The distribution shows that there is a continuum of intolerance to PTVs with a general paucity of observed versus expected truncations in the coding genome. The gray line has a slope of 1.

## Discussion

This work suggests that heterozygous protein truncating variants have greater functional consequences than generally considered. This concept is supported by the identification of a substantial proportion of genes that do not tolerate loss of one of the two gene copies, and by the evidence for a gradient of haploinsufficiency across a large proportion of the coding genome. Heterozygous PTVs are rarely compensated at the gene expression level, as shown in our previous work ^13^ and in recent analyses ^7^. Despite the absence of dosage compensation, Rivas et al. suggest that homeostatic mechanisms at the cellular level maintain biological function ^7^. However, we show clear evidence that over 10% of the genes cannot be compensated, while an additional 10 to 15% of truncated genes may be rescued by incomplete penetrance or compensatory mutations, or because the truncating variants are of limited functional impact.

The importance of these variants has also been observed in model organisms. Studies in mice show that when homozygous knockout mutants are not viable, up to 71.7% of heterozygous PTVs have phenotypic consequences ^20^. The systematic phenotyping of knockout mice also demonstrates that haploinsufficiency might be more common than generally suspected ^21^. However, a practical limitation of the above approaches, in particular in animal studies, is that observation of phenotypes resulting from damaging mutations may require exposure to specific triggers or environmental interactions ^6,21^. In contrast, in humans, life-long exposures may eventually reveal a phenotypic trait or disease associated with heterozygous gene truncations ^8^. Here, clinical symptoms could be observed later in life, and present sporadically – not necessarily within a pedigree. This is illustrated by a recent report on the consequences of haploinsufficiency of cytotoxic T-lymphocyte-associated protein 4 gene (*CTLA-4*) presenting as undiagnosed or misdiagnosed sporadic autoimmune disorder in the second to fifth decades of life ^22^. Despite the prevalence of rare heterozygous PTVs, there has been more attention to the occurrence of homozygous truncations (human knockouts). We argue that homozygous truncations result from high allele frequency variants that are less likely to carry functional consequences (the exception being recessive disorders in a population).

There are a number of possible limitations to the present study. In the modeling work, we analyzed rare variants (less than < 1% allele frequency) to focus on *de novo* events and for consistency with the *de novo* mutation rates estimated by Samocha et al. ^12^. Nevertheless our estimates held true when the analysis was restricted to singleton variants, or when we analyzed all variants irrespective of allele frequency (**Supplementary Figure 2**). We did not have primary control on sequencing coverage for some of the exome sequence datasets that could result in ascertainment errors. To correct for this potential bias, we discarded genes where the observed number of synonymous mutations deviated from expectation. The intolerance of genes to *de novo* truncation was assessed across combined human populations. Therefore, estimations of the haploinsufficient genome account for the fraction of haploinsufficient genes common to all humans. Intolerance to heterozygous PTVs should be regarded as a different concept than gene sequence conservation. PTVs in a conserved gene might have a recessive mode of inheritance and are thus potentially observable in a viable individual. On the other extreme, positively selected genes could be haploinsufficient upon heterozygous truncation. These considerations notwithstanding, we consistently identified a quantifiable fraction of the human genome that is intolerant to heterozygous PTVs, with an estimated lower bound of 9.5%.

The prevalent nature of rare heterozygous PTVs suggests that a map of “essentiality” on the basis of dominant loss of function is within reach. The concept of the essential genome has been explored in analyses of minimal bacterial genomes ^23^, mouse knockout studies ^24^, studies of transposon or chemical mutagenesis ^25^, and in studies that used CRISPR-Cas9 genome-editing technology ^26,27^. Here, we propose that mapping the haploinsufficient genome will improve the understanding of the genetic architecture of diseases. In agreement with the recent work of Li et al.,^6^ we argue that the burden of rare human heterozygous variation is an unexplored source of diversity of phenotypic traits and diseases.

## Materials and methods

**Exomes**. We collected exome data from public and non-public sources (**Supplementary Table 3**). We considered these individuals as representing the general population. Variants were filtered based on Hardy-Weinberg equilibrium (discarded if p < 1×10^−8^). For public data sets, variants were called at the data source with their respective pipelines. For non-public data sets, sequence reads were aligned using BWA, and called with Haplotypecaller using GATK 3.1. Variants were annotated with SnpEff 3.1 and filtered as described in ^28–30^. Only transcripts from autosomal protein coding genes reliably annotated by the Consensus Coding Sequence (CCDS, Release 12 04/40/2013) project^11^ that underwent the full process of CCDS curation (′Public′ status in CCDS terminology, n=17,756) were considered. As a reference background throughout all analyses, a total number of 16,521 autosomal protein coding genes was obtained by considering genes with available *de novo* mutation rate from Samocha et al. ^12^ and with at least one synonymous, missense, stop-gain or frameshift variant detected in the exome data. We discarded genes where the observed number of synonymous mutations deviated from expectation (see below). For consistency with ^12^, we only retained variants mapping within the limits of the reference transcript used to assess the *de novo* mutation rate per gene. Furthermore, only rare stop-gain and frameshift variants (allele frequency < 1%) were considered to assess the deviation from neutral expectations. Throughout the study we considered each rare variant as a single *de novo* event of mutation, irrespective of the number of individuals in which it was observed.

**Models of haploinsufficiency and noise**. Under a neutral model, the expected number of *de novo* PTVs (stop-gain or frameshift) in a gene is determined by its probability of *de novo* mutation (assessed from the sequence context and gene length) ^12^ and the number of sequenced individuals. However, potential intolerance to heterozygous truncation would decrease the expected number of *de novo* PTVs as a consequence of embryonic or early life lethality. To model the expected number of variants in a gene accounting for potential deleterious effects, we used two approaches.

First we evaluated the relative distribution of PTVs across genes (hereafter the model A). This model assumes that genes tolerating heterozygous truncation will be found truncated in the population according to their relative probability of *de novo* mutation (relative to the rest of genes), while a fraction of genes will not be observed as truncated due to early lethality. Based on the relative distribution of observed PTVs, this approach avoids issues of systematic false negative errors, though is still subject to false positive calls. Alternatively, we assessed a second model (hereafter the model B) in which the absolute number of *de novo* PTVs in a gene is estimated from the probability of *de novo* PTVs and the absolute number of observed *de novo* synonymous coding variants in that gene.

Model A is formulated as follows. In a neutral case we expect that the relative fraction of variants in a given gene is equal to 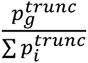, i.e. the relative distribution of observed variants follows the distributsion of *de novo* mutation rates. As some genes might harbor fewer or more mutations than the expectation, the relative model’s expected variant count for a gene g is defined as:

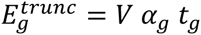

where *V* stands for the total number of observed truncating variants summed over all genes, 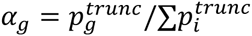, and *t*_*g*_ accounts for the gene specific deviances from the neutral case. Assuming two classes of genes (named HI for haploinsufficient and HS for non-haploinsufficient) with a class-specific *t*_*i*_ we get the following expectations for a gene g:

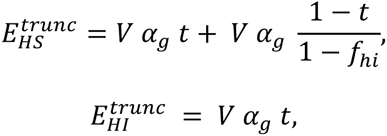

where *f*_*hi*_ is the fraction of the total number of genes that belongs to the HI class. This model distributes a fixed number of variants to all genes according to their *de novo* variation rates modulated by haploinsufficiency and the penetrance of haploinsufficient genes, and is equivalent to taking *V* samples from a multinomial distribution with α weights.

To formulate model B, we assume that the expected number of *de novo* synonymous mutations is

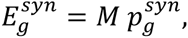

where 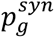 is the *de novo* rate of synonymous mutations in a gene *g* and *M* is a constant. Following ^12^ we estimate *M* from the regression of the observed number of synonymous mutations 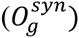 in a gene on 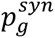:

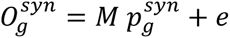

To avoid genes with low coverage, we disregarded from the analysis those genes whose residual in the above regression is higher than 3 times the standard deviation of all residuals. We note that, in contrast to ^12^ we omit the intercept term in this regression, because we expect no variants in a gene for which 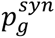 equals zero.

Having estimated *M*, the expected number of PTVs in a gene *g* is given by:

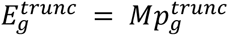

Introducing gene specific differences in the number of observed PTVs we write:

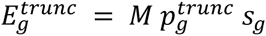

where *s*_*g*_ accounts for both gene specific differences and systematic errors. We do not estimate *s*_*g*_ for each gene, but assume that genes can be classified into two groups (haploinsufficient and non-haploinsufficient), each having a distinct class specific value of *s*.

To estimate the fraction of genes intolerant to heterozygous PTVs we use the following mixture model. We define a random variable *x*_*g*_ as the number of variants in gene *g*. A latent random variable *z*_*g*_ can take two values: *HI* or *HS* and has the probability density distribution:

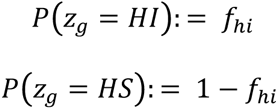

where the parameter *f*_*hi*_ represents the fraction of genes intolerant to heterozygous PTVs. The conditional probability distribution of *x*_*g*_ given *z*_*g*_ is defined as:

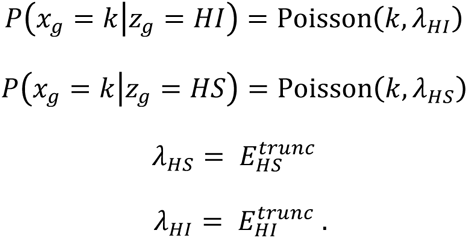

Marginalizing over the values of the latent variable *z*_*g*_ yields the probability density distribution of *x*_*g*_ as:

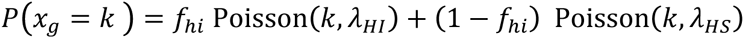

The probability that a gene acquiring *k* variants is:

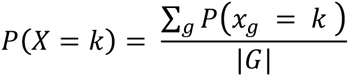

The model’s three parameters (*p*_*hi*_, *r*, *p*) are estimated by fitting the cumulative density distribution of *X* to the empirical cumulative density distribution of the data by least-squares fitting using the Nelder-Mead simplex numerical optimization algorithm (as implemented in the Apache Commons Math library). This method provided better estimates for reproducing the distribution of variant counts per gene compared to other alternatives considered (**Supplementary Figure 3**). In order to estimate the variability of the inferred model parameters we repeated the parameter estimation on 500 bootstrap replicates. Each bootstrap replicate was generated by resampling of the list of genes with replacement.

Using the estimated parameters we calculate the posterior probability of haploinsufficiency for gene *g* as:

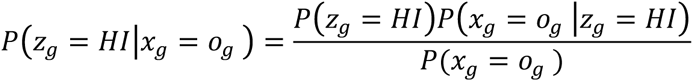

where *o*_*g*_ is the observed number of PTVs in the gene *g*.

**Characteristics of haploinsufficient genes**. Gene sets were obtained from the Reactome pathway database version 40 (http://www.reactome.org/). dN/dS values were assessed as described in ^13^. Degree of connectivity in the protein-protein interaction network was obtained from the OGEE database (http://ogeedb.embl.de/). Paralogs were counted using Ensembl Biomart’s ′Human Paralog Ensembl Gene ID′ attribute. Genes in protein complexes were obtained from Gene Ontology term GO:0043234 (named “protein complex”). Genes affecting cell viability in CRISPR-Cas9 experiments were collected from ^26,27^. Severity of protein truncation was assessed by the NutVar score (http://nutvar.labtelenti.org) ^13^. For the assessment of depletion or enrichment of functional gene sets we used one tailed hypergeometric test. We adjusted the p-values by the Benjamini- Hochberg method to correct for multiple testing. We tested pathways with at least 100 elements only.

## Acknowledgments

The authors would like to thank the NHLBI GO Exome Sequencing Project and its ongoing studies; The 1000 Genomes Project, the TwinsUK Cohort; The Avon Longitudinal Study of Parents and Children; The Genome of the Netherlands Project; The Swiss HIV Cohort Study and The National Institute of Environmental Health Science Environmental Genome Project. We are also grateful for access to exome sequence data from the CoLaus cohort, which was sequenced as part of a partnership between the Wellcome Trust Sanger Institute, the CoLaus principal investigators and the Quantitative Sciences dept. of GlaxoSmithKline. We acknowledge the helpful comments of Viktor Müller. Part of the computations were performed at the Vital-IT (http://www.vital-it.ch) Center for high-performance computing of the SIB Swiss Institute of Bioinformatics.

## Supplementary materials

**Table S1:**
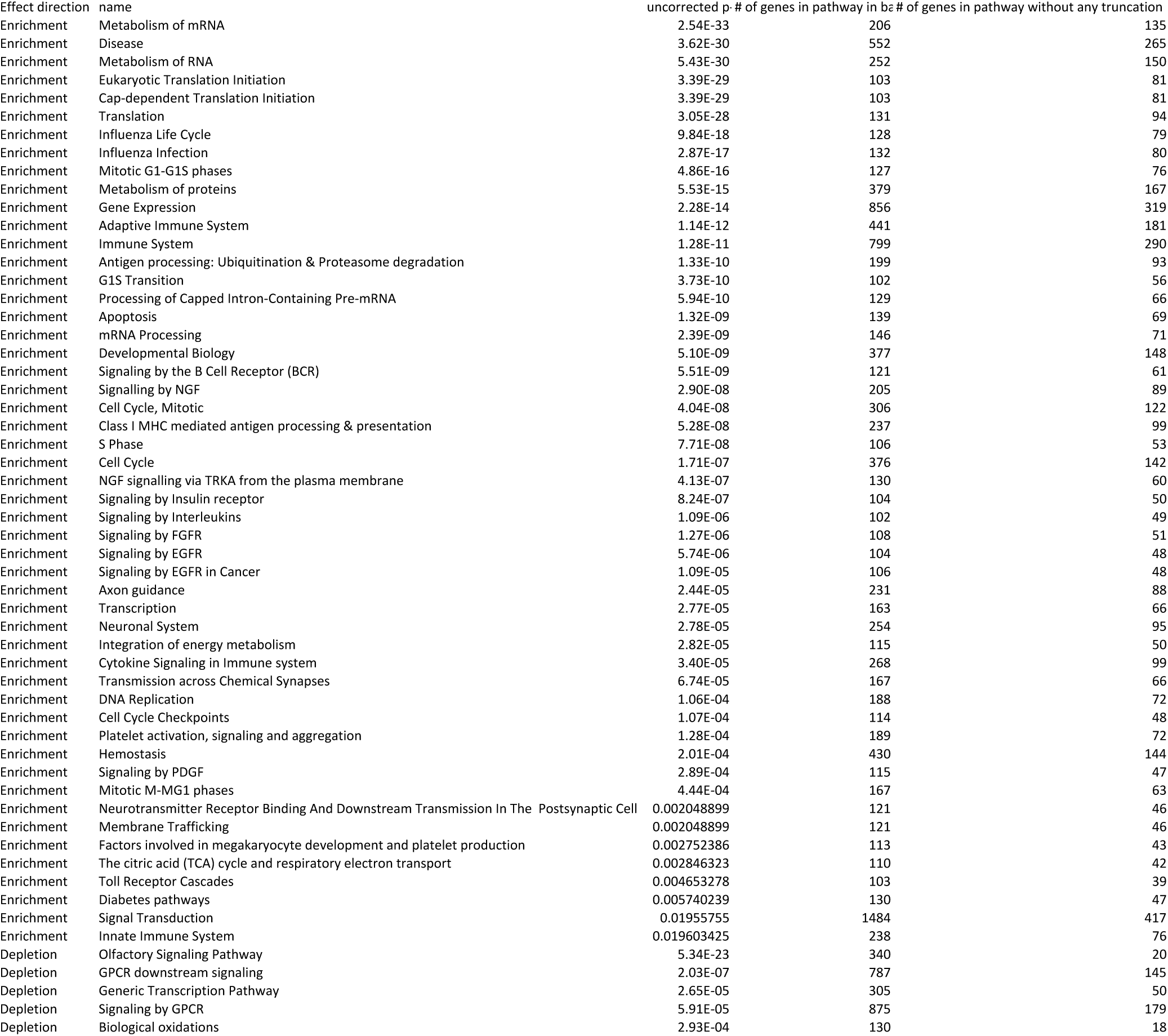
Enrichment tests results against Reactome pathways for genes without PTVs. Only significant results are shown as judged by 5% FDR calculated using the Benjamini-Hochberg procedure.

**Table S2:**
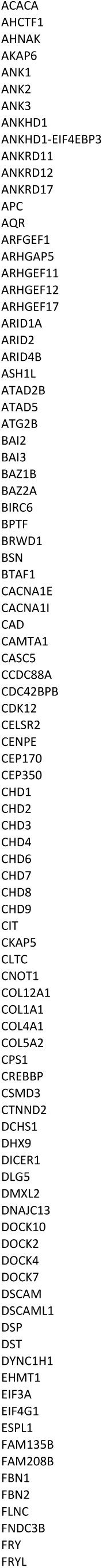
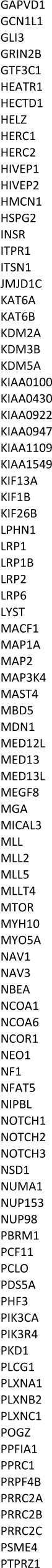
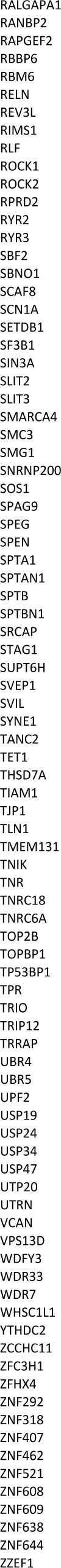
Genes with higher than 0.99 posterior probability of being intolerant to heterozygous PTVs.

**Table S3:** Data sources.

| Project name | Sample size | Reference | URL |
| --- | --- | --- | --- |
| NHLBI Exome Sequencing Project | 6502 | (34) | http://evs.gs.washington.edu/EVS/ |
| UK10K | 2432 | (35) | http://www.bristol.ac.uk/alspac/ |
| 1000 Genomes Project | 1092 | (36) | http://www.1000genomes.org/ |
| Genome of the Netherlands | 498 | (37) | http://www.nlgenome.nl |
| Cohort Lausanne | 426 | (38) | http://www.colaus.ch/ |
| Swiss HIV Cohort Study | 500 | (39) | http://www.shcs.ch/ |
| NIEHS Environmental Genome Project | 95 | (40) | http://evs.gs.washington.edu/niehsExome |
| Genome of J. Craig Venter | 1 | (41) | http://huref.jcvi.org/ |
| Total | 11546 |  |  |

**Figure S1:**
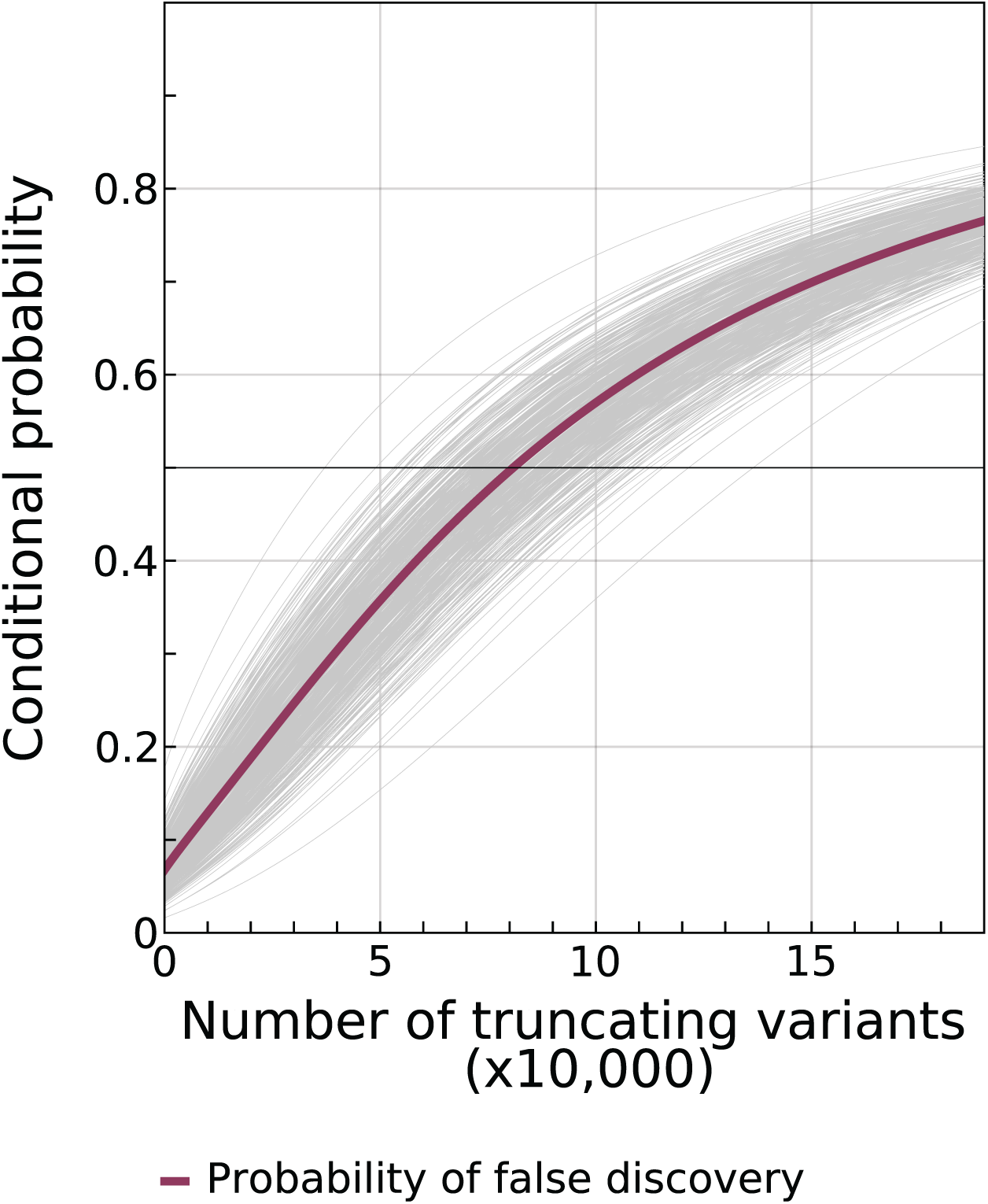
Conditional probability that when observing a gene truncated for the first time, the gene is intolerant to PTVs. When the conditional probability crosses 50% (at 90,000 PTVs) biological and technical noise become the main source of truncations. We estimate that 40,000 exomes are required to sample 90,000 PTVs using the jackknife projection as in^31^.

**Figure S2:**
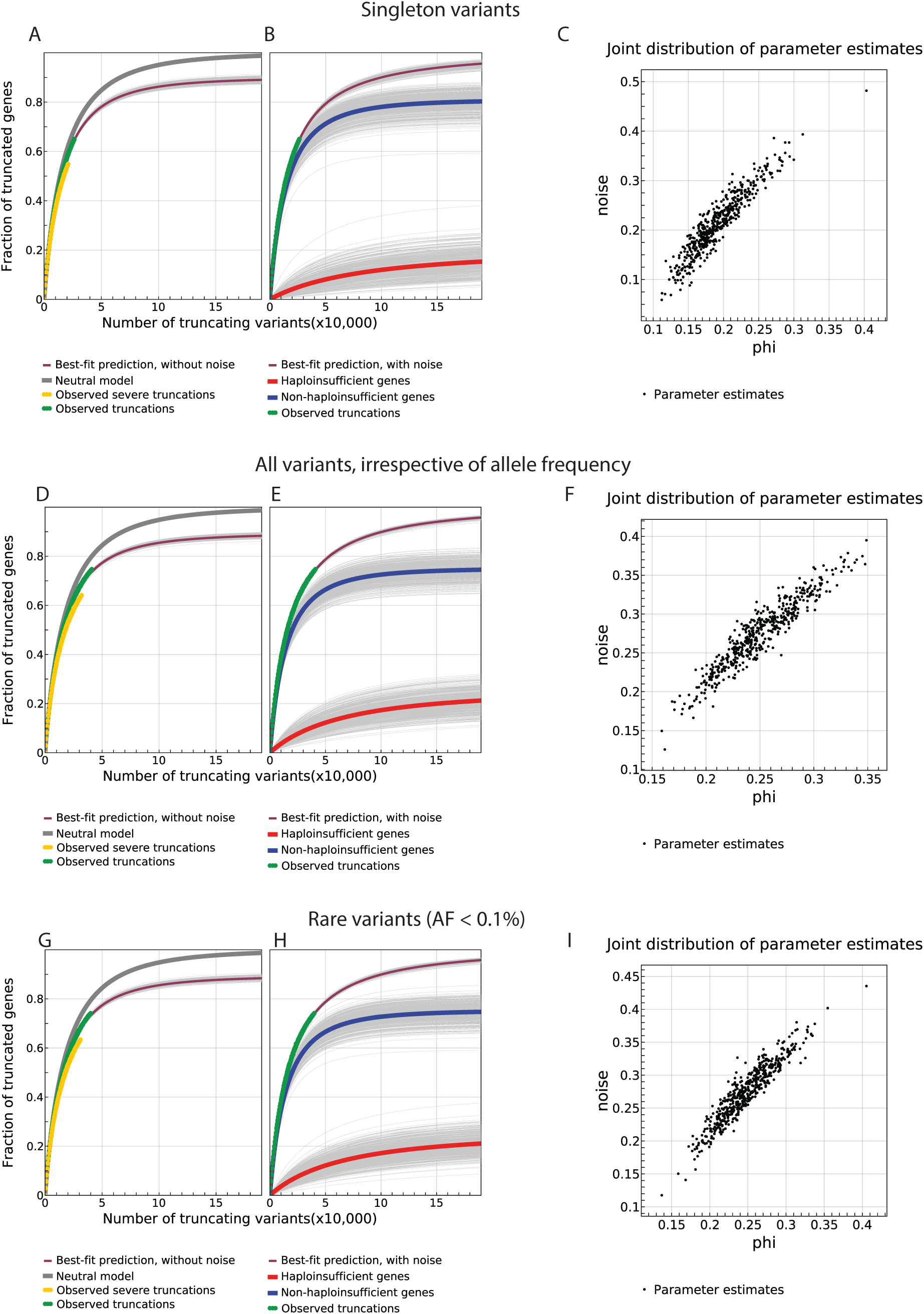
Distribution of parameter estimates and predictions of the model A. Analysis considers only singletons (A-C), all variants irrespective of allele frequency (D-F) or rare variants (G-I).

**Figure S3:**
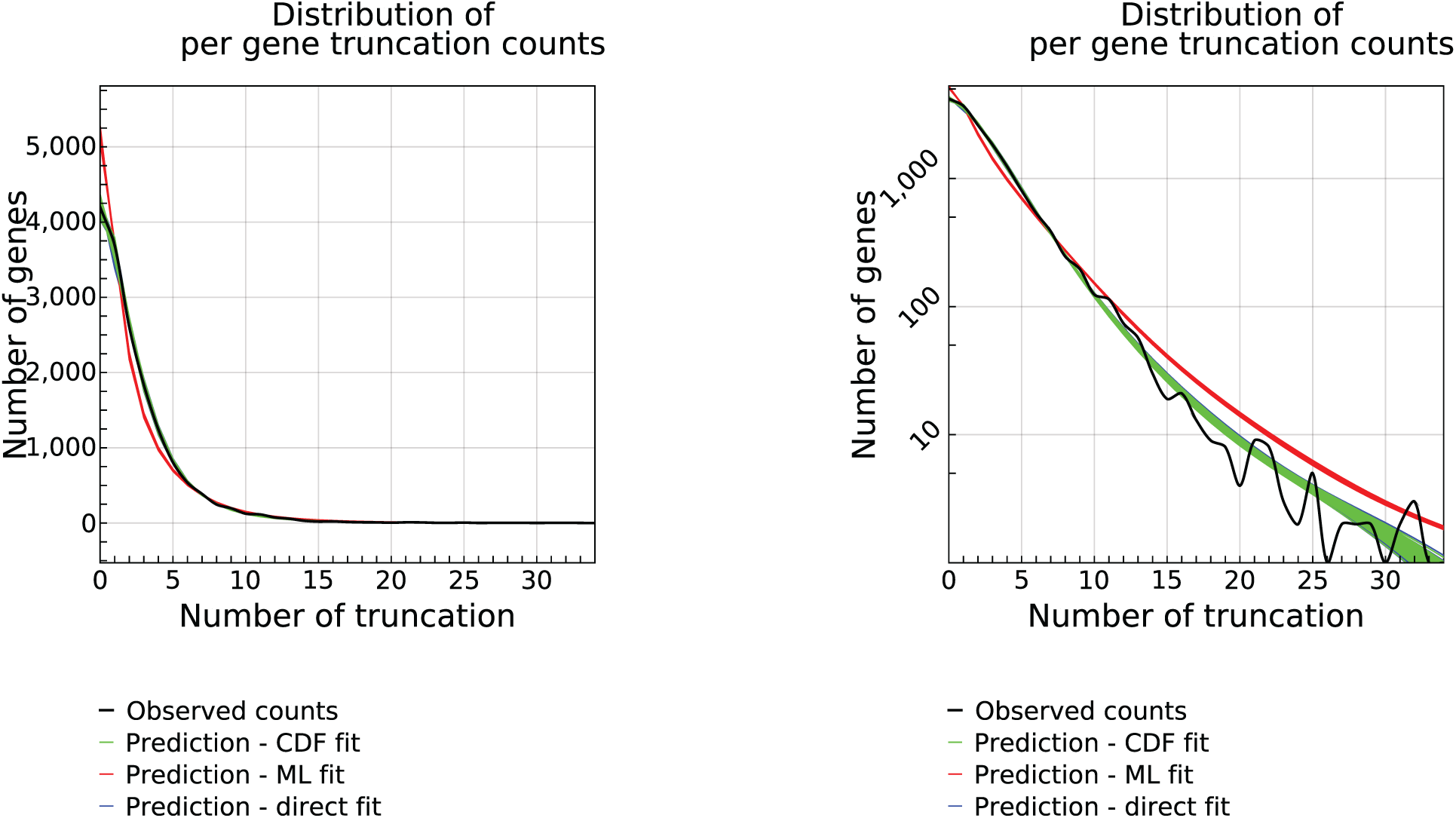
Distribution of variant counts per gene as observed or predicted under best-fit parameters of model A using 3 different estimation techniques. A: linear space, **B:** log space. Black curve: observed counts, red curve: prediction based on least-squares fit to the cumulative distribution function (see Online Methods), green curve: maximum likelihood estimate, blue curve: least squares fit to the accumulation curve of truncated genes as shown in Figure 1. The CDF method was chosen and maximum likelihood was discarded because its estimates did not fit the observations.

## References and Notes

1. Tennessen, J.A. et al. Evolution and functional impact of rare coding variation from deep sequencing of human exomes. Science 337, 64–9 (2012).

2. Nelson, M.R. et al. An abundance of rare functional variants in 202 drug target genes sequenced in 14,002 people. Science 337, 100–4 (2012).

3. MacArthur, D.G. et al. Guidelines for investigating causality of sequence variants in human disease. Nature 508, 469–76 (2014).

4. MacArthur, D.G. et al. A systematic survey of loss-of-function variants in human protein-coding genes. Science 335, 823–8 (2012).

5. Sulem, P. et al. Identification of a large set of rare complete human knockouts. Nat Genet 47, 448–52 (2015).

6. Li, A.H. et al. Analysis of loss-of-function variants and 20 risk factor phenotypes in 8,554 individuals identifies loci influencing chronic disease. Nat Genet (2015).

7. Rivas, M.A. et al. Human genomics. Effect of predicted protein-truncating genetic variants on the human transcriptome. Science 348, 666–9 (2015).

8. Lim, E.T. et al. Distribution and medical impact of loss-of-function variants in the Finnish founder population. PLoS Genet 10, e1004494 (2014).

9. Fisher, E. & Scambler, P. Human haploinsufficiency-one for sorrow, two for joy. Nat Genet 7, 5–7 (1994).

10. Huang, N., Lee, I., Marcotte, E.M. & Hurles, M.E. Characterising and predicting haploinsufficiency in the human genome. PLoS Genet 6, e1001154 (2010).

11. Pruitt, K.D. et al. The consensus coding sequence (CCDS) project: Identifying a common protein-coding gene set for the human and mouse genomes. Genome Res 19, 1316–23 (2009).

12. Samocha, K.E. et al. A framework for the interpretation of de novo mutation in human disease. Nat Genet 46, 944–50 (2014).

13. Rausell, A. et al. Analysis of stop-gain and frameshift variants in human innate immunity genes. PLoS Comput Biol 10, e1003757 (2014).

14. Dang, V.T., Kassahn, K.S., Marcos, A.E. & Ragan, M.A. Identification of human haploinsufficient genes and their genomic proximity to segmental duplications. Eur J Hum Genet 16, 1350–7 (2008).

15. Rieux-Laucat, F. & Casanova, J.L. Immunology. Autoimmunity by haploinsufficiency. Science 345, 1560–1 (2014).

16. Szamecz, B. et al. The genomic landscape of compensatory evolution. PLoS Biol 12, e1001935 (2014).

17. Wall, J.D. et al. Estimating genotype error rates from high-coverage next-generation sequence data. Genome Res 24, 1734–9 (2014).

18. Fang, H. et al. Reducing INDEL calling errors in whole genome and exome sequencing data. Genome Med 6, 89 (2014).

19. Weisenfeld, N.I. et al. Comprehensive variation discovery in single human genomes. Nat Genet 46, 1350–5 (2014).

20. Ayadi, A. et al. Mouse large-scale phenotyping initiatives: overview of the European Mouse Disease Clinic (EUMODIC) and of the Wellcome Trust Sanger Institute Mouse Genetics Project. Mamm Genome 23, 600–10 (2012).

21. White, J.K. et al. Genome-wide generation and systematic phenotyping of knockout mice reveals new roles for many genes. Cell 154, 452–64 (2013).

22. Kuehn, H.S. et al. Immune dysregulation in human subjects with heterozygous germline mutations in CTLA4. Science 345, 1623–7 (2014).

23. Hutchison, C.A. et al. Global transposon mutagenesis and a minimal Mycoplasma genome. Science 286, 2165–9 (1999).

24. Bradley, A. et al. The mammalian gene function resource: the International Knockout Mouse Consortium. Mamm Genome 23, 580–6 (2012).

25. Venken, K.J. & Bellen, H.J. Chemical mutagens, transposons, and transgenes to interrogate gene function in Drosophila melanogaster. Methods 68, 15–28 (2014).

26. Shalem, O. et al. Genome-scale CRISPR-Cas9 knockout screening in human cells. Science 343, 84–7 (2014).

27. Wang, T., Wei, J.J., Sabatini, D.M. & Lander, E.S. Genetic screens in human cells using the CRISPR-Cas9 system. Science 343, 80–4 (2014).

28. McKenna, A. et al. The Genome Analysis Toolkit: a MapReduce framework for analyzing next-generation DNA sequencing data. Genome Res 20, 1297–303 (2010).

29. Li, H. & Durbin, R. Fast and accurate long-read alignment with Burrows-Wheeler transform. Bioinformatics 26, 589–95 (2010).

30. Reumers, J. et al. SNPeffect: a database mapping molecular phenotypic effects of human non-synonymous coding SNPs. Nucleic Acids Res 33, D527-32 (2005).

31. Gravel, S. et al. Demographic history and rare allele sharing among human populations. Proc Natl Acad Sci U S A 108, 11983–8 (2011).

32. Petrovski, S., Wang, Q., Heinzen, E.L., Allen, A.S. & Goldstein, D.B. Genic intolerance to functional variation and the interpretation of personal genomes. PLoS Genet 9, e1003709 (2013).

## References

[34] Exome Variant Server, NHLBI GO Exome Sequencing Project.

[35] A. Boyd, et al., International journal of epidemiology 42, 111 (2013).

[36] G. R. Abecasis, et al., Nature 491, 56(2012).

[37] The Genome of the Netherlands Consortium, T. Genome, Nature genetics 46, 818 (2014).

[38] M. Firmann, et al., BMC cardiovascular disorders 8, 6 (2008).

[39] J. Fellay, et al., PLoS genetics 5, e1000791 (2009).

[40] NIEHS Environmental Genome Project, Seattle, WA.

[41] S. Levy, et al., PLoS Biology 5, 2113 (2007).

